# Truncated power-law distribution of group sizes in antelope

**DOI:** 10.1101/411199

**Authors:** Pranav Minasandra, Kavita Isvaran

**Affiliations:** Undergraduate programme, Indian Institute of Science, Bengaluru, India; Centre for Ecological Sciences, Indian Institute of Science, Bengaluru, India

## Abstract

Group size distributions are instrumental in understanding group behaviour in animal populations. We analysed group size data of the blackbuck, *Antilope cervicapra,* from six different field sites to estimate the group size distribution of this antelope. We show that an exponentially truncated power law (called the polylog distribution in this paper) is the best fitting distribution, against the simple power law and lognormal distributions as other contenders, and the exponential distribution as a control. To show this, we use two likelihood based methods (AICs and likelihood ratios). Finally, we show that polylog distribution parameters can be used to better understand group dynamics, by using them to explore how habitat openness affects group behaviour.

## Introduction

Animals show a spectacular variety of social grouping patterns (Krause & Ruxton, 2002). Group sizes vary widely among species. Within species, group sizes may vary across populations and even at small spatial and temporal scales within a population. For example, many species display fission-fusion groups, that is, groups that form by the splitting of larger groups or the merging of smaller groups. Groups in such populations are dynamic and vary in size through the day. Most studies on group size variation and group dynamics focus on the trade-offs underlying grouping patterns and measure the costs and benefits to individuals from being a part of a group of a particular size (Alexander, 1974; Pulliam & Caraco, 1984). Studies of mechanisms underlying the behaviour of social groups examine how individuals of a group move in space (Ballerini et al, 2008). Larger-scale studies have examined how ecological conditions may affect trade-offs and thereby grouping patterns in a population (Isvaran, 2007). Such studies represent grouping patterns in a population using measures, such as mean and Typical Group Size (Jarman 1974). Thus studies of adaptiveness of grouping and mechanisms associated with grouping, have typically focussed either on individuals in groups of particular sizes or on mean/typical group sizes in a population. Rarely do studies of grouping patterns attempt to understand the overall distribution of group sizes in a population (Bonabeau and Dagorn, 1995; Griesser et al, 2011).

Understanding group size distributions in wild populations on the field can help us in testing models of group dynamics. This can be done by answering the question: “Does the model we propose predict the group size distribution observed in nature?”. It is difficult to obtain substantial data involving group dynamics in the field. For example, for fission-fusion groups, the exact rules involved in the merging and splitting of groups is unknown, and many possible rules have been reviewed in Pays et al, (2007). Group decisions on merging and splitting are very complex functions of numerous variables, such as group size, spatial location of members, duration of time the group has been in existence, etc. Measuring these variables is difficult, and it is relatively easier to collect data on group sizes and find their distribution. If this distribution does not match the one predicted by a model, it would indicate that the model needs improvement.

A way to fully understand group size distributions is to find the Probability Mass Function (PMF) of the group size, f(*s*), where:

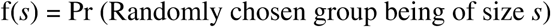

Group sizes can take integer values ≥ 1. Therefore, we are looking for a PMF f:N → [0, 1] where N is the set of Natural numbers. Several such distributions have been proposed. Okubo (1986) proposed that group sizes are exponentially distributed, based on maximising the entropy of the group size distribution. However, an unstated assumption in his model is that the group size distribution has a finite expectation. This is not necessarily true. Bonabeau et al (1999), working with a mechanistic model where two groups merge whenever they meet, derive that group sizes are power-law distributed (Table 1: zeta distribution), with an exponent of 1.5. Such a distribution has an infinite expectation. Intuitively, there must be a truncation to this heavy tailed power-law, because there are a finite number of organisms in a population, and because groups also split at some rate. This leads to the truncated power law. (Table 1: polylog distribution)

**Table 1:** Group Size Distributions and their PMFs. *s* can take natural number values, except in the case of the lognormal distribution, where *s* can take all positive real values. We can calculate the expectation (distribution-mean), distribution-variance, etc., using these PMFs

| Distribution | PMF |
| --- | --- |
| Exponential | $(1 - e^{-\lambda})e^{-\lambda(s-1)}$ |
| Zeta | $s^{-b} / \zeta(b)$ |
| Lognormal | $\exp(-(\ln s - \mu)^2 / 2 \sigma^2) / s \sigma (2\pi)^{0.5}$ |
| Polylog | $(s^{-b} e^{-\lambda s}) / \text{Li}_b(e^{-\lambda})$ |

A power-law or truncated power-law distribution allows the frequent occurrence of very large numbers (in this case, group sizes), and is often seen in the literature (Klaus et al, 2011; Clauset et al, 2007; Clauset et al, 2009; Griesser et al, 2011). Both these ‘heavy-tailed’ distributions often carry large variances (sometimes infinite variance with the simple power-law). As the terms ‘power-law’ and ‘truncated power-law’ are ambiguous, we propose to use more formal names for the distributions involved. In this paper, we refer to the power-law as the zeta distribution, and the exponentially truncated power-law as the polylog distribution (These names are based on the normalising coefficients of these distributions). The polylog distribution is the zeta distribution with an exponential cutoff. Its PMF is given in Table 1, where Li_*b*_ is the logarithmic integral of order *b*, also called the polylog function.

We examine group size distributions in the model antelope *Antilope cervicapra*, the Blackbuck. Blackbuck occur in open grasslands, and form groups containing members of all ages and sexes. Blackbuck groups follow fission-fusion dynamics, with their group sizes changing in the span of a single day (Mungall, 1978). Groups sizes are distributed with a high variance (Ranjitsinh, 1989).

In this paper, we have contrasted all the PMFs listed in Table 1 against actual data collected from six wild populations of blackbuck in different years between 1998 and 2001, and different geographical locations across the range of blackbuck in India (Table 2). We use simple likelihood-based data analysis tools to compare the different models. To demonstrate the usefulness of analysing group size distributions, we test the effect of habitat openness on grouping. For this, we investigate how group size distribution parameters vary with habitat openness, and make interpretations accordingly.

**Table 2:** Information about the six datasets. used in this study. Data were collected as described in the Data collection subsection. Location and major habitat type have been reproduced from Isvaran (2005)

| Geographical location | Number of groups | Duration of sampling | Location | Major habitat type |
| --- | --- | --- | --- | --- |
| Velavadar | 701 | Jan 2000 - May 2000 | 21°56' N,<br>72°10' E | Grassland, shrubland, mudflats |
| Rollapadu | 109 | August 1998 | 15°52' N,<br>78°18' E | Grassland |
| Point calimere | 103 | October 1998 | 10°18' N,<br>79°51' E | Forest with grassy openings |
| Rehekuri | 86 | October 1999 | 19°42' N,<br>75°44' E | Forest, grassland |
| Nannaj | 80 | October - November 1999 | 17°41' N,<br>75°56' E | Grassland |
| Tal Chhapar | 52 | October 1998 | 27°88' N,<br>74°58' E | Grassland, shrubland |

## Methods

### Data collection

The blackbuck antelope, native to the Indian subcontinent, is found in a wide range of habitats, from dry grasslands to open woodlands (Jhala and Isvaran, 2016). They are group-living grazers. Group sizes vary widely both between and within populations. Grouping, in this species, appears to reflect a balance between the benefits of avoiding predation and the costs of increased competition for food. Average group sizes appear to increase with ecological conditions associated with predation-related benefits and decrease with those associated with food-competition (Isvaran, 2007).

The data we use here were collected from six different blackbuck populations (Table 2). At 5 of the 6 populations, regular censuses were conducted on foot during which the study area was systematically covered. For every group sighted, the number of animals and the sex and stage (fawn, juvenile, adult) were recorded. In one population, 1000 x 100 m strip transects were used to record group sizes, since the habitat was too dense for census methods (details of total counts and transects in Isvaran (2007)). Apart from social groups, male blackbuck defend mating territories from where they display to females and attempt to mate with receptive females that visit their territories (Mungall, 1978; Ranjitsinh, 1989; Jhala and Isvaran, 2016). Territorial males could be identified through their behaviour and were excluded from the analyses of social group size distribution (Isvaran, 2007).

### Analysis

For each population, we fitted all the distributions to the data, where the best possible fit was found using Maximum Likelihood (ML) Estimation. We used Akaike Information Criteria (AICs) and Log-of-Likelihood-Ratios to evaluate the relative fits. We extensively used the python library powerlaw written by Alsott et al (2014)^3^ throughout the analysis.

ML estimates were arrived at as described in (Clauset et al, 2009) and were computed using the code by (Alsott et al, 2014).

AICs were calculated for each dataset for each distribution. The AIC is a statistic that is used to quantify model optimality, by factoring in how well the model explains the data, as well as the complexity of the model. It penalises overfitting, and also enables us to compare various models. It is computed using the formula

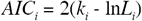

where *k*_*i*_ is the number of parameters and *L*_*i*_ is the likelihood of the data under the *i*^th^ model. The AIC always assumes a positive value. The model with the lowest AIC is assumed to be the best model.

We see in the results (Table 3) that the AICs follow the same trend across all six populations. The AIC values, with a few minor deviations, suggest that the following order of distributions (from better to worse fit) always holds:

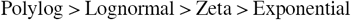

**Table 3:** AICs and (δAICs) for all datasets for all the distributions under consideration. The smallest AIC is in bold. Notice that, AICs almost always decrease as we go down the rows. This tells us that the polylog distribution is the best fit to the data.

| Distribution | Population |  |  |  |  |  |
| --- | --- | --- | --- | --- | --- | --- |
|  | Velavadar | Rollapadu | Point Calimere | Rehekuri | Nannaj | Tal Chhapar |
| Exponential | 5566.31<br>(1111.01) | 711.52<br>(88.21) | 588.71<br>(39.58) | 480.30<br>(7.31) | 639.48<br>(62.51) | 475.75<br>(100.9) |
| Zeta | 4525.83<br>(70.53) | 637.35<br>(14.04) | 569.44<br>(20.31) | 502.28<br>(29.21) | 598.14<br>(21.27) | 378.96<br>(4.11) |
| Lognormal | 4468.80<br>(13.5) | 628.55<br>(5.24) | **549.13**<br>(0) | 476.42<br>(3.43) | 582.76<br>(5.79) | 378.36<br>(3.51) |
| Polylog | **4455.30**<br>(0) | **623.31**<br>(0) | 549.22<br>(0.09) | **472.99**<br>(0) | **576.97**<br>(0) | **374.85**<br>(0) |

To further verify this result, we use another likelihood based method, the log-of-likelihood-ratio (LLR) method. This method provides a way to directly compare distributions pairwise. The LLR is defined as

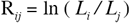

where *L*_*i*_ and *L*_*j*_ are the likelihoods of the data assuming model *i* and *j* respectively.

Due to the trend observed above, we compared only the following 3 pairs: (a) Exponential and Zeta; (b) Zeta and Lognormal; and (c) Lognormal and Polylog. As described in (Vuong, 1989) and (Clauset et al, 2009), we have reported the normalised LLR (which we have denoted by R instead of R_norm_ for the sake of convenience), and the corresponding p-value.

Additionally, to explore the meaning of *b* and *λ*, we examined the relation between the estimated parameters and habitat openness. Habitat openness was measured at the same time as the group size,s and the values used are from Isvaran (2005)

Finally, to find the performance of frequently reported statistics, we investigated the performance of the mean, typical group size, and also the MLE parameters of *b* and λ through the simulations described in Appendix II.

## Results

The MLE estimated parameter values for all distributions from Table 1, for all the datasets, are reported in Appendix I.

Across all six datasets, AICs for the polylog distribution were always the lowest, except in the Point Calimere data set, where the lognormal distribution has a negligibly lower AIC than the polylog (Table 3). Therefore, the polylog distribution described blackbuck group data sets very well. The results of the LLR test (Table 4) closely agree with those shown by computing the AIC values. This confirms that the Polylog distribution best describes Blackbuck group data (Figure 1). Finally, we found an increasing trend in *b* and 1/*λ* with habitat openness (Figure 2). (Spearman rank correlation test, and the obtained *ρ* values are *b* vs habitat openness: *ρ* = 0.927, *ρ* = 0.007; and *λ* vs habitat openness: *ρ* = -0.927, *ρ* = 0.007)

**Table 4:** LLR values with their associated *p*-values for the distributions. The results here perfectly match those shown by the AIC values in Table 3.

| Distributions compared |  | Velavadar | Rollapadu | Point Calimere | Rehekuri | Nannaj | Tal Chhapar |
| --- | --- | --- | --- | --- | --- | --- | --- |
| **Zeta vs Exponential** | R | 8.91 | 3.26 | 0.76 | -1.55 | 1.79 | 4.37 |
| | $p$ | $4.91 \times 10^{-19}$ | 0.0011 | 0.4469 | 0.1207 | 0.0719 | $1.22 \times 10^{-5}$ |
| **Lognormal vs Zeta** | R | 6.56 | 3.28 | 3.25 | 3.69 | 2.94 | 2.71 |
| | $p$ | $5.15 \times 10^{-11}$ | 0.0010 | 0.0011 | 0.0002 | 0.0032 | 0.0067 |
| **Polylog vs Lognormal** | R | 3.15 | 3.37 | -0.03 | 1.16 | 2.81 | 1.98 |
| | $p$ | 0.0016 | 0.0007 | 0.9699 | 0.2463 | 0.0049 | 0.0047 |

**Figure 1:**
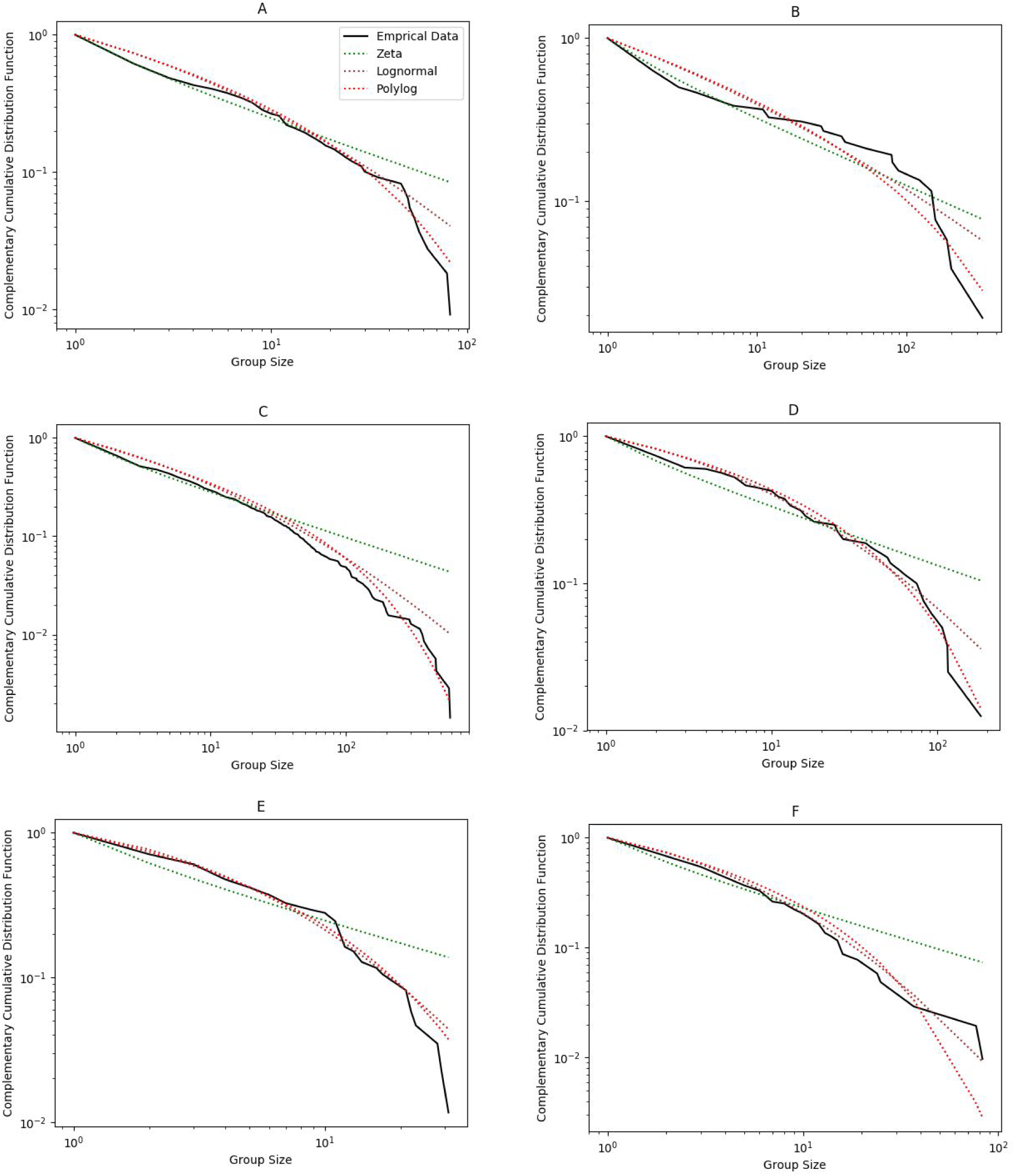
Complementary Cumulative distribution functions of the fits of the distributions in Table 1 to the datasets. (except exponential, whose performance was extremely poor (Table 3 and Table 4)) and of the data. (A) Rollapadu, (B) Tal Chhapar, (C) Velavadar, (D) Nannaj, (E) Rehekuri, and (F) Point Calimere. The polylog distribution has the best fit in all cases.

**Figure 2:**
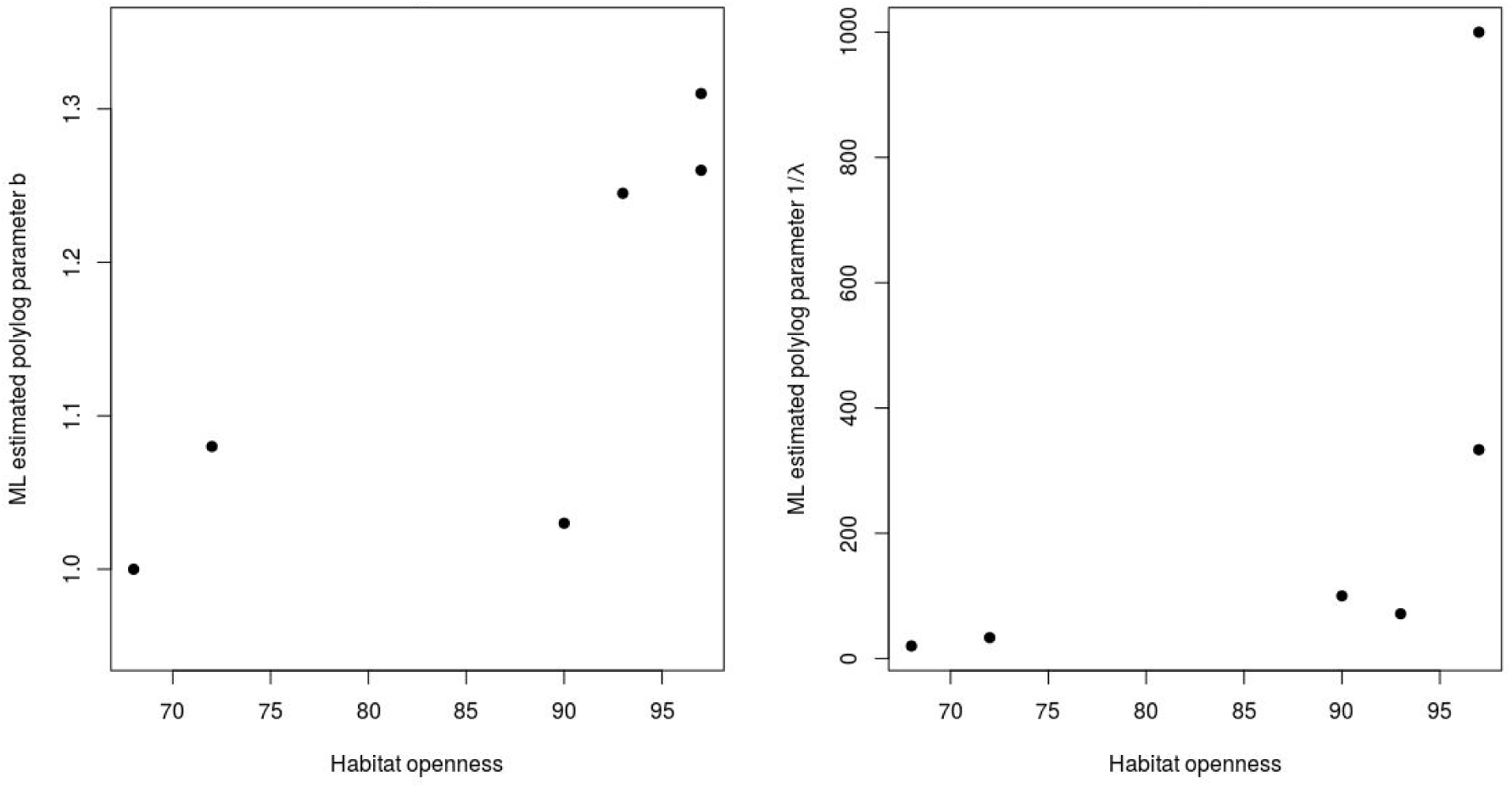
Polylog parameters *b* and 1/*λ* vs habitat openness. Clearly, both these parameters increase with increasing habitat openness. 1/*λ* is plotted because it is more intuitive, it can be interpreted as the group size where the power-law → exponential transition occurs. Correlations of *b* and *λ* with H.O are significant, as reported in Results.

## Discussion

We compared the fit of four plausible distributions to group size data collected from six wild blackbuck populations. Our results indicate that group size distributions in blackbuck typically follow the polylog distribution and its parameters can provide insights into group dynamics and responses to ecological conditions.

The polylog distribution was the best fit for the observed group sizes in all datasets. The only place where the order of fits was not clear was in the Point Calimere dataset, and in this case the lognormal performed almost as well as the polylog distribution. The polylog distribution was followed by the lognormal, the zeta, and the exponential.

These datasets have been collected from populations separated by hundreds of kilometres and several years (Table 2). This suggests that the results will apply across all populations of blackbuck. As blackbuck groups show typical fission-fusion behaviour, the same results will possibly apply across species that follow fission-fusion dynamics.

The polylog parameters, *b* and *λ* have biologically relevant meanings, and can be used to derive insights about group dynamics. Lower values of *b* and *λ* imply an increased probability of large groups occurring. In Bonabeau et al (1999) *b* and *λ* are interpreted as follows: *b* captures the dimensionality of the environment, where it increases with increasing freedom of motion, reaching a maximum value of 1.5 when there is no constraint on movement (i.e, increasing freedom of movement decreases the probabilities of groups meeting, and so results in smaller groups in general). *λ* reflects the probability of splitting of a group. Higher the *λ*, the more the groups tend to split. Moreover, the finiteness of the population size also constrains the value of *λ*. The values we find for *b* and *λ* (Appendix I) also agree well with those predicted in Bonabeau et al (1999). Further, in case of the polylog fit, the values of *b* estimated from our datasets tend to be close to 1, which is essentially the ‘logarithmic distribution’ used in Griesser et al (2011).

We also show that the estimated polylog parameters *b* and *λ* can be used to make ecological inferences. Examining variation in habitat openness with *b* and *λ* across populations, we found that *b* increases with habitat openness. This can be explained using the interpretation of the parameters in Bonabeau et al (1999). The more open the habitat becomes, the less often groups meet each other due to there being fewer constraints on movement. 1/*λ* also increased (i.e *λ* decreased) with increasing habitat openness. This leads us to make the prediction that groups split less often in more open habitats. One explanation for this is that, in open habitats, individuals in any group can see all other members with more ease than in a less open habitat (Gerard et al, 2002). Due to this, cohesiveness of groups is likely to increase in more open habitats.

Our results also have implications for testing models of group dynamics. Given our results, any model of group dynamics that predicts a polylog or zeta stationary group size distribution might be the appropriate model for group dynamics for blackbuck and other species with similar fission-fusion behaviour. Both predictions are correct because the polylog distribution is simply the zeta distribution truncated due to the splitting of groups (polylog distribution → zeta distribution as *λ* → 0).

Mean and Typical Group Size in any heavy-tailed distribution are expected to carry large variances. Further, different pairs of *b* and *λ* can have the same value for these statistics. Our simulations (Appendix II; Fig A.2, A.3) confirm that sample means carry a large variance in the polylog distribution. Hence, they become difficult to use in ecological considerations. Therefore, reporting ML estimated *b* and *λ* as statistics of group sizes in animals following fission-fusion dynamics, alongside conventionally reported statistics can be informative. We find that *b* and *λ* are estimated with a bias (Appendix II; Fig A.4). However, the advantage in reporting these parameters is that, these values remove ambiguity in the data, as *b* and *λ* comprehensively capture the group size distribution. As estimates *b* and *λ* are biased with a consistent sign (Appendix II), reporting them is still potentially useful.

To conclude, our data supports an exponentially truncated power law group size distribution in wild populations. This is in concordance with the predictions of the model by Bonabeau et al (1995, 1999), and also with other field-based observations, e.g. Griesser et al (2011). The extensiveness of our data, coupled with the agreement of results across six geographically distant populations, points towards the probable generality of this distribution of group sizes across all species that display similar behaviours in grouping.

## Acknowledgements

We are very grateful to Vishwesha Guttal and Manjunath Krishnapur for useful discussions. We thank Aditi Ajith Pujar, Dhanya Bharath, and Vibhuti Shastri for their valuable comments on the manuscript.

### Appendix I

#### Maximum likelihood estimated values of parameters

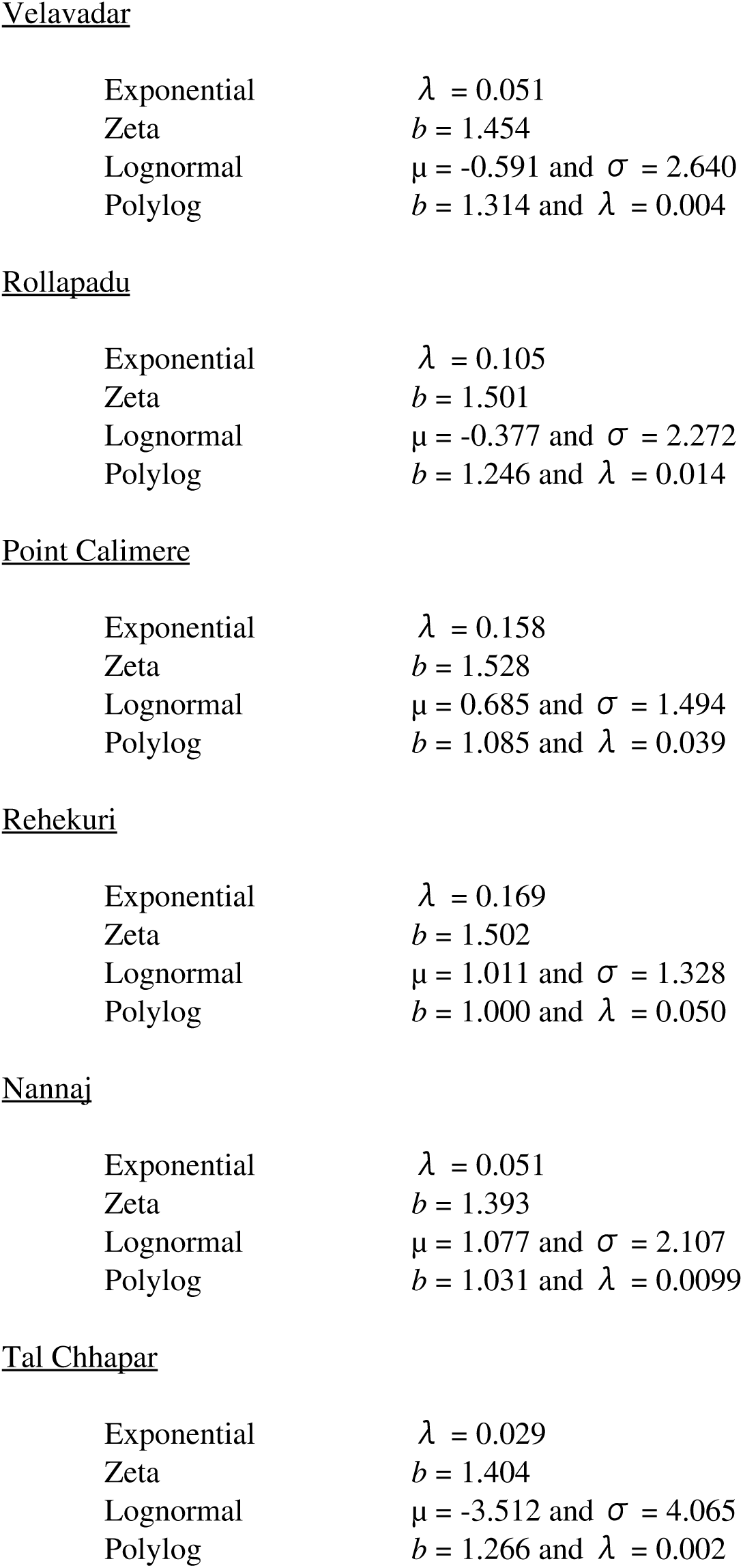

Velavadar
|  |  |
| --- | --- |
| Exponential | $\lambda = 0.051$ |
| Zeta | $b = 1.454$ |
| Lognormal | $\mu = -0.591$ and $\sigma = 2.640$ |
| Polylog | $b = 1.314$ and $\lambda = 0.004$ |
| Rollapadu |  |
| Exponential | $\lambda = 0.105$ |
| Zeta | $b = 1.501$ |
| Lognormal | $\mu = -0.377$ and $\sigma = 2.272$ |
| Polylog | $b = 1.246$ and $\lambda = 0.014$ |
| Point Calimere |  |
| Exponential | $\lambda = 0.158$ |
| Zeta | $b = 1.528$ |
| Lognormal | $\mu = 0.685$ and $\sigma = 1.494$ |
| Polylog | $b = 1.085$ and $\lambda = 0.039$ |
| Rehekuri |  |
| Exponential | $\lambda = 0.169$ |
| Zeta | $b = 1.502$ |
| Lognormal | $\mu = 1.011$ and $\sigma = 1.328$ |
| Polylog | $b = 1.000$ and $\lambda = 0.050$ |
| Nannaj |  |
| Exponential | $\lambda = 0.051$ |
| Zeta | $b = 1.393$ |
| Lognormal | $\mu = 1.077$ and $\sigma = 2.107$ |
| Polylog | $b = 1.031$ and $\lambda = 0.0099$ |
| Tal Chhapar |  |
| Exponential | $\lambda = 0.029$ |
| Zeta | $b = 1.404$ |
| Lognormal | $\mu = -3.512$ and $\sigma = 4.065$ |
| Polylog | $b = 1.266$ and $\lambda = 0.002$ |

### Appendix II

#### Performance of distribution parameters, mean, and typical group size

We investigated the behaviour of two statistics, sample mean and typical group size, which are often reported in the literature. We assume that group sizes follow a polylog distribution. We also evaluated the performance of the ML estimators of the parameters of the polylog distribution.

For this, we generated 500 datasets, each containing 100 data points, for each pair of *b* and *λ* shown in Fig A.1. The sample size of 100 was used keeping in mind ecological data. For every dataset, we calculated sample mean, Typical Group Size (TGS), and the ML estimated values of *b* and *λ*. Further, for every *b*-*λ* pair, we found the expected values of the mean and the typical group size according to Equations A1 and A2.

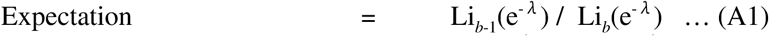

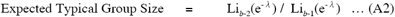

Fig A.2 and A.3 represent the performance of the mean and TGS. The values of these estimated from the data are associated with a large variance. This property needs to be kept in mind when reporting these statistics for ecological data.

**Figure A.1:**
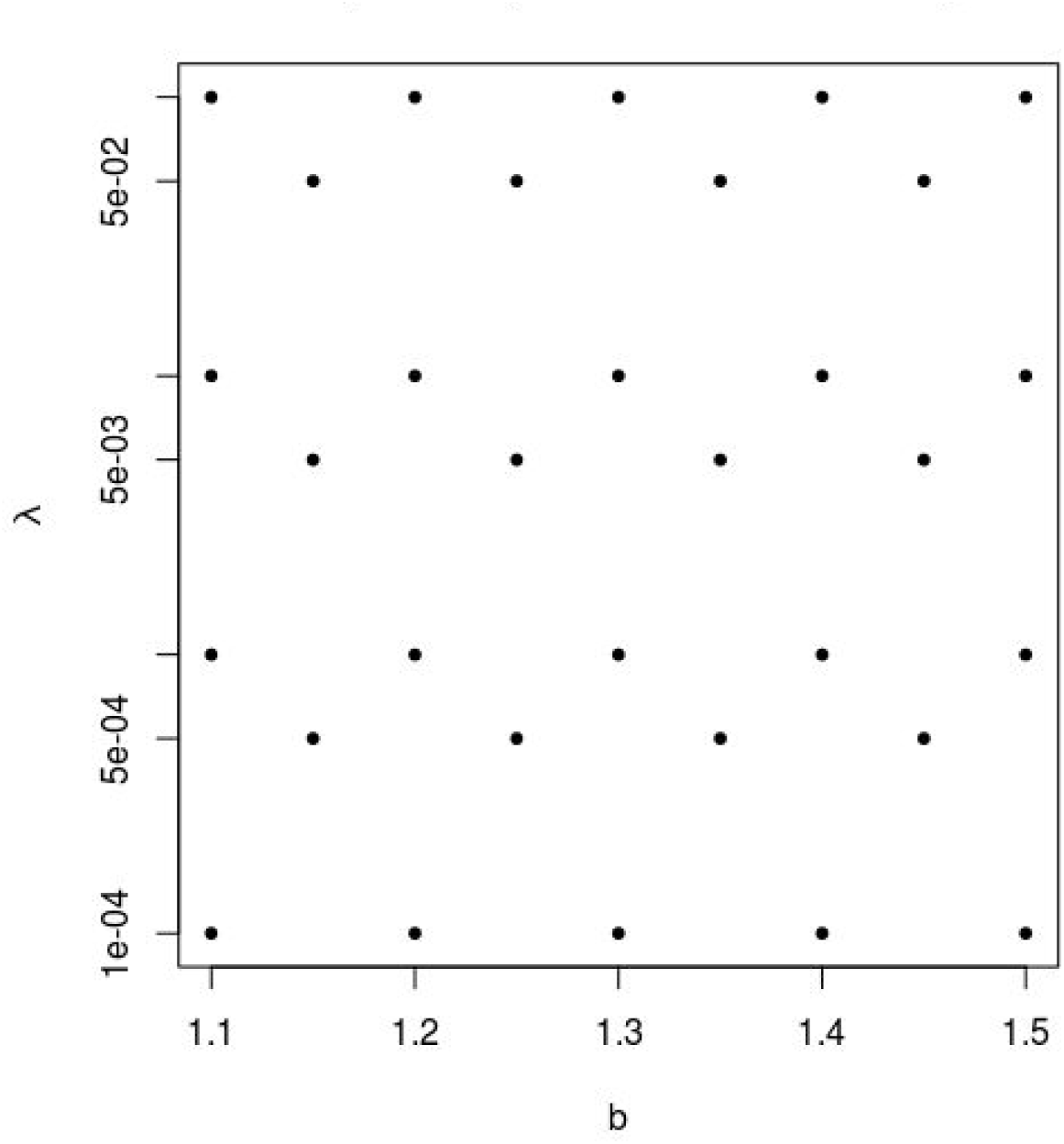
Each dot represents a pair of *b* and *λ* on which the analyses mentioned in Appendix II were performed.

**Figure A.2:**
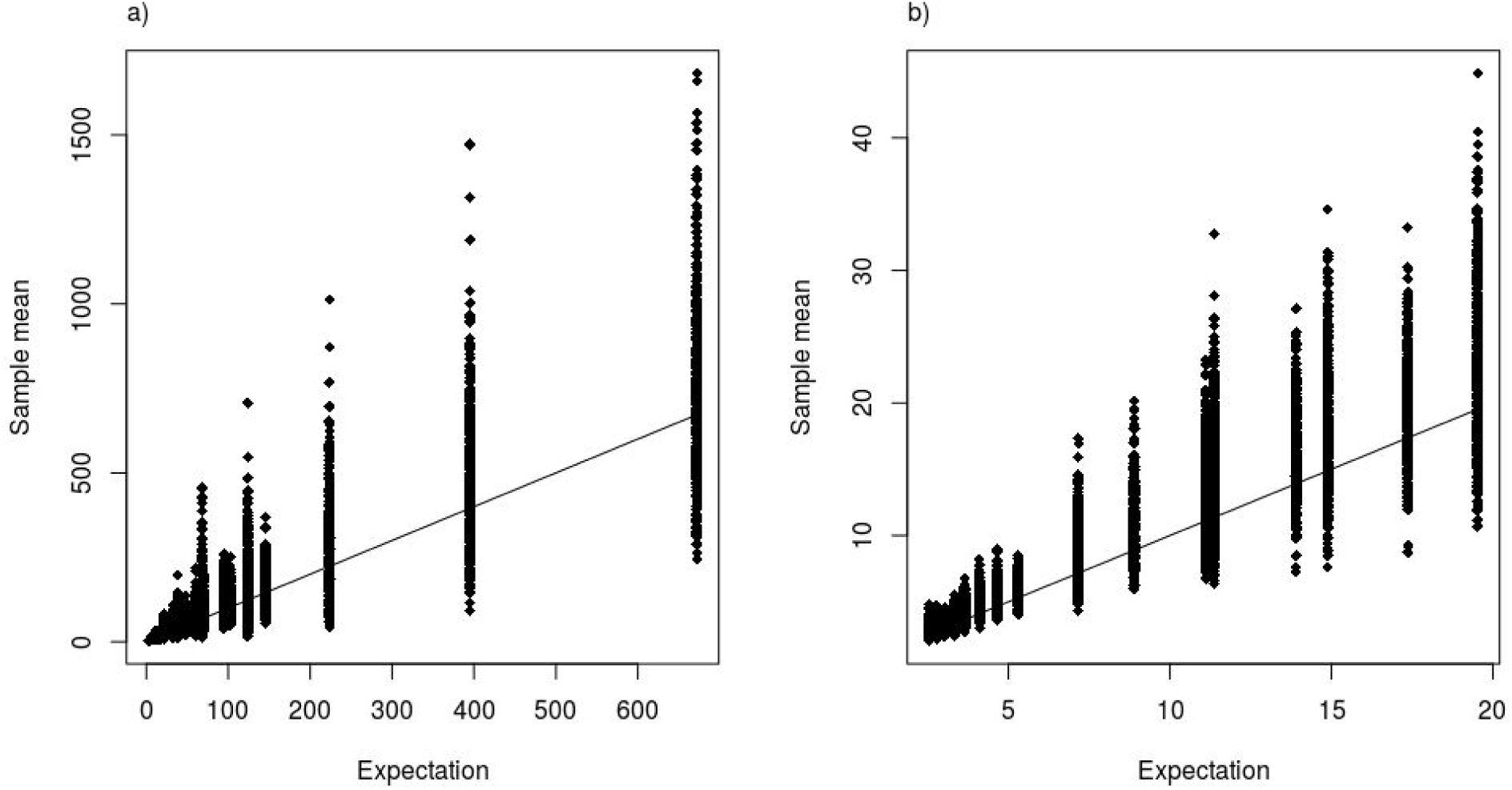
Performance of the sample mean (N = 100 numbers per dataset); a) over a large interval, and b) over a biologically relevant interval, as seen from our datasets. The line in both figures is the *y* = *x* line. Each point in these graphs corresponds to a dataset containing 100 data points.

**Figure A.3:**
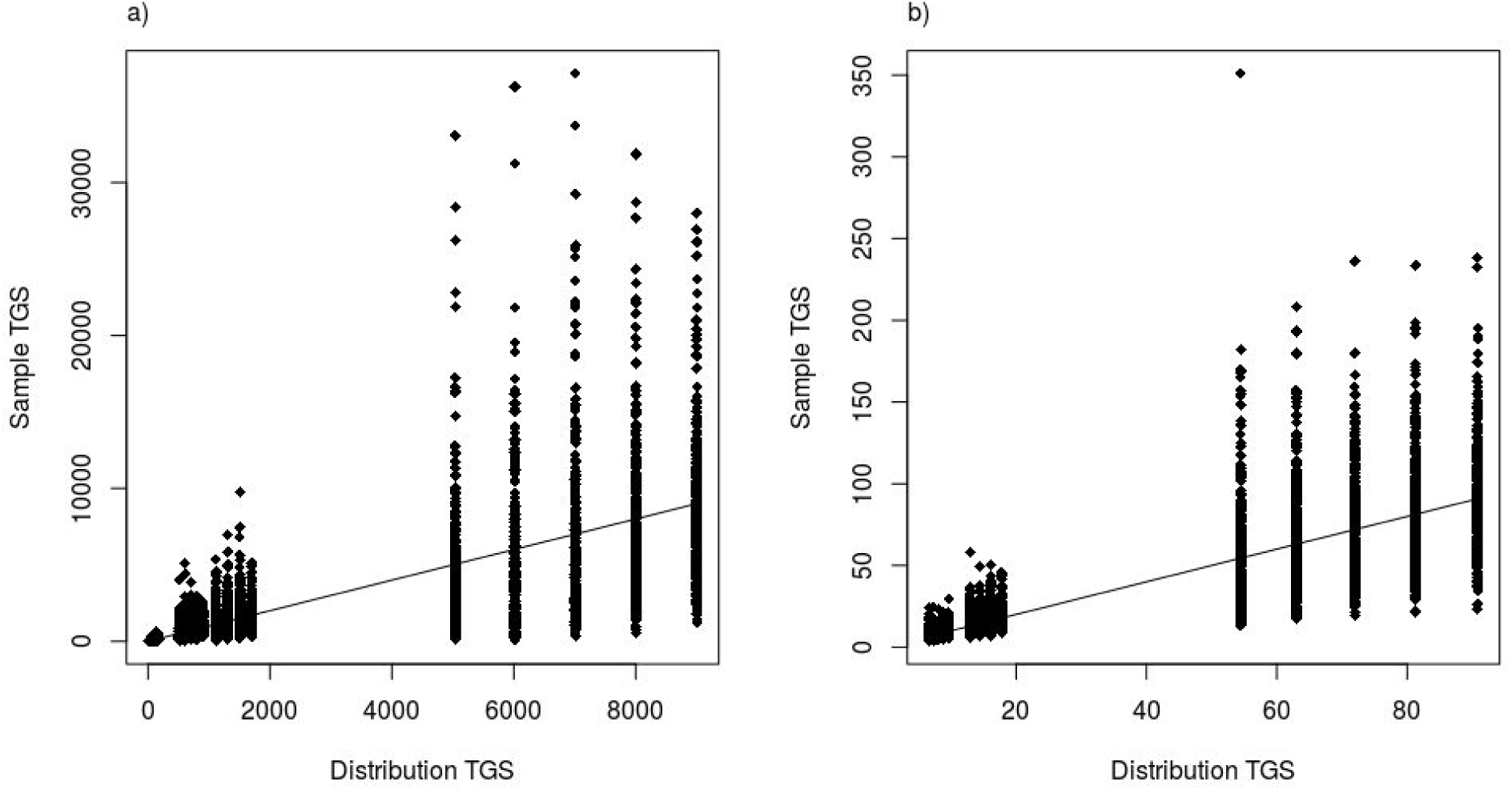
Performance of the Typical Group Size (N = 100 numbers per dataset); a) over a large interval, and b) over a biologically relevant interval, as seen from our datasets. The line in both figures is the *y* = *x* line. Each point in these graphs corresponds to a dataset containing 100 data points.

Further, we found a negative bias in the estimator for *b* and a positive bias in the estimator for *λ*. To quantify bias, we use a law of large numbers approach: Bias is defined as the expectation of the deviation of the estimate from the true value. We approximate the expectation with the sample mean of deviations across 500 datasets per parameter pair.

The biases so calculated increased in magnitude with increasing *b* and *λ*. Fig A.3 shows the effects of bias over *b*-*λ* space. Table A.T1 provides the values of bias at the points analysed.

**Figure A.3:**
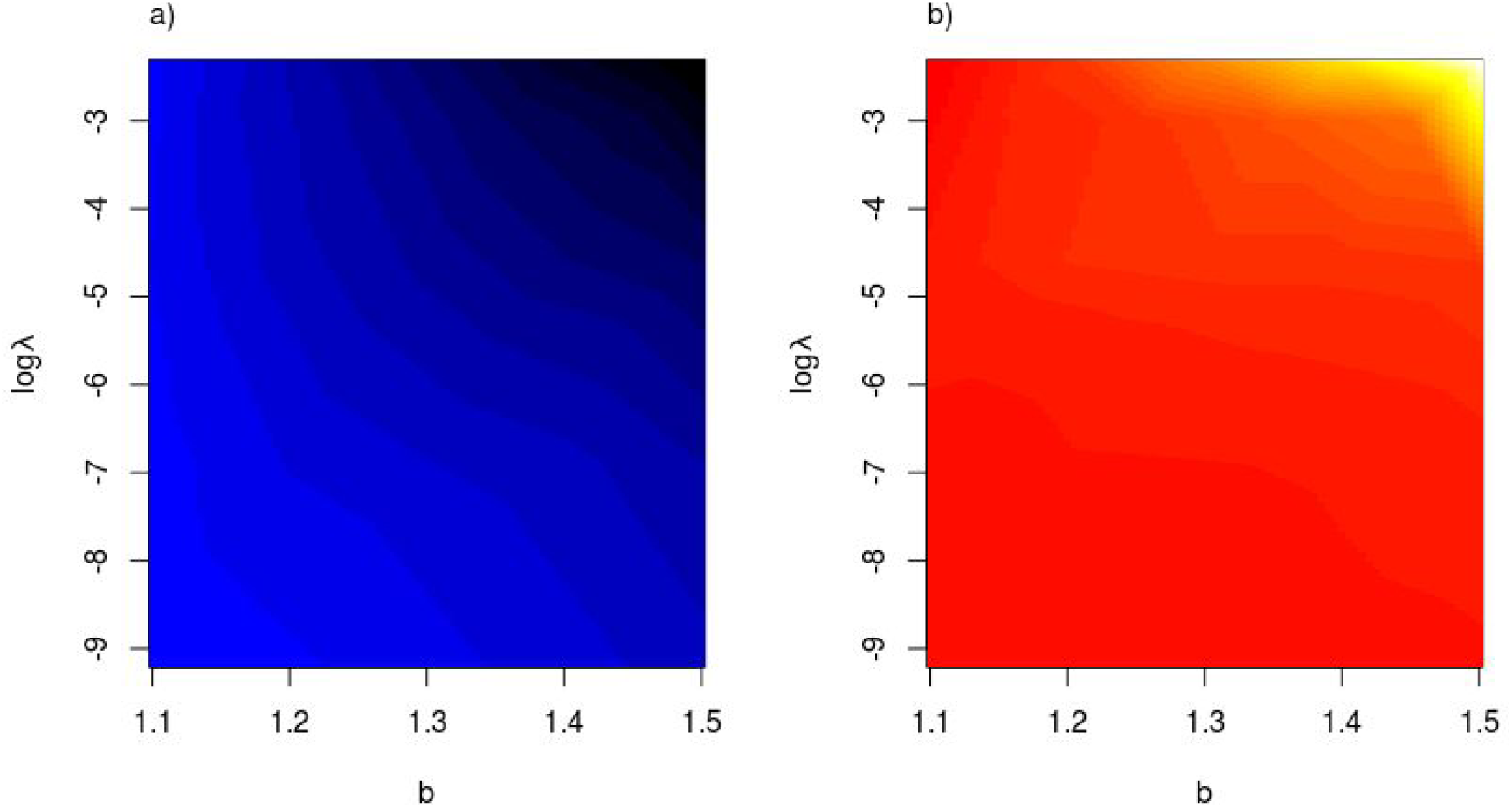
Biases in the parameters of the polylog distribution; a) Bias in estimating *b*, and b) Bias in estimating *λ*. The bias values are provided in Table A.T1, and these have been interpolated using the R package by Akima et al (2013) to make this graph. Bias in *b* is negative, whereas in *λ* it is positive. Both biases increase in magnitude with increasing *b* and *λ*. In a) Blue → Black is increasingly negative bias, whereas in b), Red → White is increasingly positive bias.

**Table A.T1:**
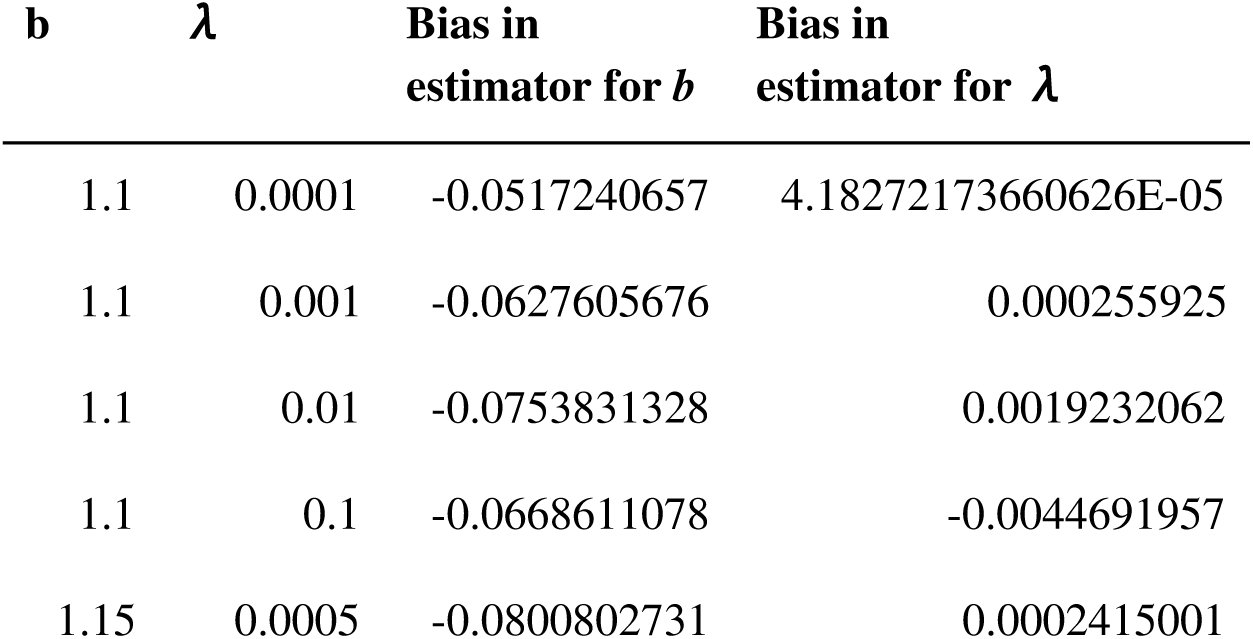
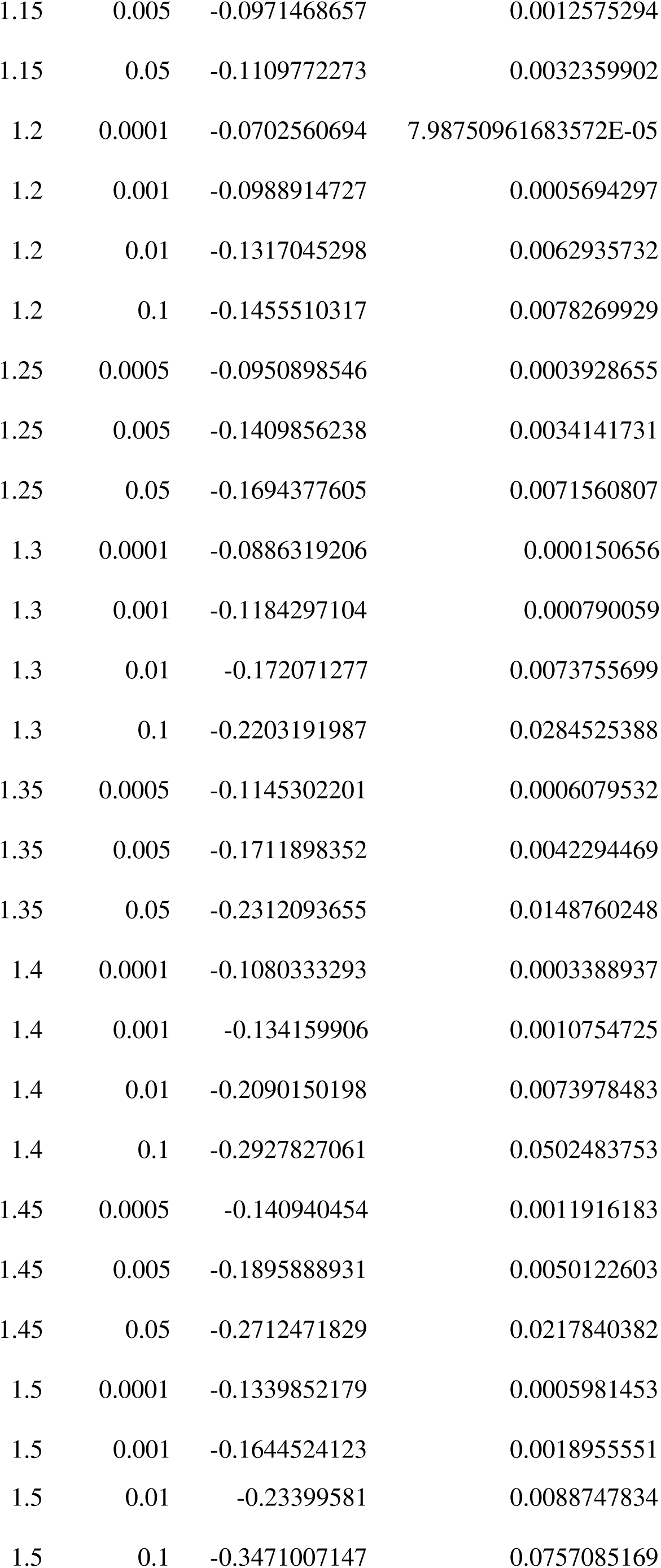
Biases in the estimators for *b* and *λ* as functions of *b* and *λ*. Each ‘bias’ in the third and fourth column represents the deviation of the estimate from the true value, averaged across 500 datasets.

| $b$ | $\lambda$ | Bias in<br>estimator for $b$ | Bias in<br>estimator for $\lambda$ |
| --- | --- | --- | --- |
| 1.1 | 0.0001 | -0.0517240657 | 4.18272173660626E-05 |
| 1.1 | 0.001 | -0.0627605676 | 0.000255925 |
| 1.1 | 0.01 | -0.0753831328 | 0.0019232062 |
| 1.1 | 0.1 | -0.0668611078 | -0.0044691957 |
| 1.15 | 0.0005 | -0.0800802731 | 0.0002415001 |
| 1.15 | 0.005 | -0.0971468657 | 0.0012575294 |
| 1.15 | 0.05 | -0.1109772273 | 0.0032359902 |
| 1.2 | 0.0001 | -0.0702560694 | 7.98750961683572E-05 |
| 1.2 | 0.001 | -0.0988914727 | 0.0005694297 |
| 1.2 | 0.01 | -0.1317045298 | 0.0062935732 |
| 1.2 | 0.1 | -0.1455510317 | 0.0078269929 |
| 1.25 | 0.0005 | -0.0950898546 | 0.0003928655 |
| 1.25 | 0.005 | -0.1409856238 | 0.0034141731 |
| 1.25 | 0.05 | -0.1694377605 | 0.0071560807 |
| 1.3 | 0.0001 | -0.0886319206 | 0.000150656 |
| 1.3 | 0.001 | -0.1184297104 | 0.000790059 |
| 1.3 | 0.01 | -0.172071277 | 0.0073755699 |
| 1.3 | 0.1 | -0.2203191987 | 0.0284525388 |
| 1.35 | 0.0005 | -0.1145302201 | 0.0006079532 |
| 1.35 | 0.005 | -0.1711898352 | 0.0042294469 |
| 1.35 | 0.05 | -0.2312093655 | 0.0148760248 |
| 1.4 | 0.0001 | -0.1080333293 | 0.0003388937 |
| 1.4 | 0.001 | -0.134159906 | 0.0010754725 |
| 1.4 | 0.01 | -0.2090150198 | 0.0073978483 |
| 1.4 | 0.1 | -0.2927827061 | 0.0502483753 |
| 1.45 | 0.0005 | -0.140940454 | 0.0011916183 |
| 1.45 | 0.005 | -0.1895888931 | 0.0050122603 |
| 1.45 | 0.05 | -0.2712471829 | 0.0217840382 |
| 1.5 | 0.0001 | -0.1339852179 | 0.0005981453 |
| 1.5 | 0.001 | -0.1644524123 | 0.0018955551 |
| 1.5 | 0.01 | -0.23399581 | 0.0088747834 |
| 1.5 | 0.1 | -0.3471007147 | 0.0757085169 |

Mean and TGS are estimated with a large variance, deviating considerably from their true values. MLE parameters *b* and *λ* are estimated with biases. However, the biases are consistent in sign over the entire biologically valid range of values. Due to this, *b* and *λ* can still be used in statistical analyses.

## Footnotes

3 Code required to perform our analysis is freely available at <u>github.com/pminasandra/distribution-analyse</u>

## References

1. Akima, H., Gebhardt, A., Petzoldt, T., & Maechler, M. (2013). akima: Interpolation of irregularly spaced data. R package version 0.5-11, URL http://CRAN.R-project.org/package=akima.

2. Alsott, J., Bullmore, E., & Plenz, D. (2014). powerlaw: a Python package for analysis of heavy-tailed distributions. PloS one, 91), e85777.

3. Alexander, R. D. (1974). The evolution of social behavior. Annual review of ecology and systematics, 5(1), 325-383.

4. Ballerini, M., Cabibbo, N., Candelier, R., Cavagna, A., Cisbani, E., Giardina, I. … & Zdravkovic, V. (2008). Empirical investigation of starling flocks: a benchmark study in collective animal behaviour. Animal behaviour, 76(1), 201–215.

5. Bonabeau, E., & Dagorn, L. (1995). Possible universality in the size distribution of fish schools. Physical Review E, 51(6), R5220.

6. Bonabeau, E., Dagorn, L., & Freon, Pierre. (1999). Scaling in animal group-size distributions. Proceedings of the National Academy of Sciences, 96(8), 4472–4477.

7. Clauset, A., Shalizi, C. R., & Newman, M. E. (2009). Power-law distributions in empirical data. SIAM review, 51(4), 661–703.

8. Clauset, A., Young, M., & Gleditsch, K. S. (2007). On the frequency of severe terrorist events. Journal of Conflict Resolution, 51(1), 58–87.

9. Gerard, J. F., Bideau, E., Maublanc, M. L., Loisel, P., & Marchal, C. (2002). Herd size in large herbivores: encoded in the individual or emergent?. The Biological Bulletin, 202(3), 275–282.

10. Griesser, M., Ma, Q., Webber, S., Bowgen, K., & Sumpter, D. J. (2011). Understanding animal group-size distributions. PLoS One, 6(8), e23438.

11. Isvaran, K. (2005). Female grouping best predicts lekking in blackbuck (Antilope cervicapra). Behavioral Ecology and Sociobiology, 57(3), 283–294.

12. Isvaran, K. (2007). Intraspecific variation in group size in the blackbuck antelope: the roles of habitat structure and forage at different spatial scales. Oecologia, 154(2), 435–444.

13. Jarman, P. (1974). The social organisation of antelope in relation to their ecology. Behaviour, 48(1), 215–267.

14. Jhala, Y. V., & Isvaran, K. (2016). Behavioural Ecology of a Grassland Antelope, the Blackbuck Antilope cervicapra: Linking Habitat, Ecology and Behaviour. In The Ecology of Large Herbivores in South and Southeast Asia (pp. 151-176). Springer, Dordrecht.

15. Klaus, A., Yu, S., & Plenz, D. (2011). Statistical analyses support power law distributions found in neuronal avalanches. PloS one, 6(5), e19779.

16. Krause, J., Ruxton, G. D., & Ruxton, G. D. (2002). Living in groups. Oxford University Press.

17. Mungall, E. C. (1978). The Indian blackbuck antelope: a Texas view(No. QL737. M86 1978.).

18. Mungall, E. C. (1998). Bucks in the black: India vs. Texas. Exotic Wildl, 8, 1–3.

19. Okubo, A. (1986). Dynamical aspects of animal grouping: swarms, schools, flocks, and herds. Advances in biophysics, 22, 1–94.

20. Pays, O., Benhamou, S., Helder, R., & Gerard, J. F. (2007). The dynamics of group formation in large mammalian herbivores: an analysis in the European roe deer. Animal Behaviour, 74(5), 1429–1441.

21. Pulliam, H. R., & Caraco, T. (1984). Living in groups: is there and optimal group size? -In: ‘Behavioural Ecology: An Evolutionary Approach’. 2nd edn.(Eds JR Krebs and ND Davies.) pp. 122–147.

22. Ranjitsinh, M. K. (1989). Indian blackbuck. Natraj Publishers.

23. Vuong, Q. H. (1989). Likelihood ratio tests for model selection and non-nested hypotheses. Econometrica: Journal of the Econometric Society, 307–333.

